# Modeling of Non-invasive Cell Membrane Potential Measurement: LTSpice Simulation and Machine Learning Analysis

**DOI:** 10.1101/2024.09.05.611360

**Authors:** Xiaofeng Ma

## Abstract

This paper presents a novel simulation approach using electric pulses to measure cell membrane potential. The primary objective was to develop a non-invasive method for accurately assessing membrane potential without altering the cell membrane or its internal components. Traditional techniques, such as voltage-sensitive dyes, often require cell incubation, which can affect membrane properties and reduce measurement accuracy. To overcome these limitations, we modeled the cell as two series-connected capacitors with a negative potential between them. By manipulating this negative potential and analyzing the charge and discharge characteristics, simulations conducted with LTspice demonstrated the feasibility of predicting membrane potential based on these characteristics.

We tested 202 groups with various capacitor combinations and measured four key parameters related to charge and discharge currents: maximum current, minimum current, total charge time, and total discharge time. Using the XGBRegressor model, we achieved a strong fit with an R^2^ score of 0.9. This indicates a robust correlation between the measured charge and discharge characteristics and the internal potential of the cell. Our findings suggest that accurate measurement of cell membrane potential is possible by attaching an electrode to the cell without disrupting membrane integrity. Thus, this simulation-based approach offers a promising and non-invasive alternative for measuring cell membrane potential.

## Introduction

Accurately measuring cell membrane potential in a non-invasive manner is crucial for various biological research areas, including health, disease, and drug discovery. The membrane potential is vital for numerous physiological processes, such as the propagation of nerve impulses, muscle contraction, and the regulation of cardiac rhythm (1,2). Abnormalities in membrane potential can lead to various health issues, including epilepsy, cardiac arrhythmias, and muscle disorders (3-5). Additionally, understanding membrane potential dynamics is essential for developing new drugs, as many pharmacological agents target ion channels and membrane receptors involved in maintaining these potentials (6).

Sharp electrodes are currently the gold standard for this measurement, but they come with drawbacks such as the risk of introducing internal solution into the cell and limitations in recording multiple cells efficiently. Previous research has made advancements towards non-invasive measurements by utilizing voltage-sensitive dyes (7-9) and micro-electrochemical techniques (10). However, these approaches may still modify cell membrane properties, emphasizing the need for a method that minimizes cell damage while accurately assessing potential.

In this study, we explored an alternative approach by attaching an electrode to the cell with high membrane resistance, creating a cell-attached configuration. By considering a system composed of two series-connected capacitors with a negative charge in between, we modeled this configuration’s behavior during charge and discharge properties when pulses are applied. This allowed us to estimate the negative charge through measurement, offering valuable insights and confirming the viability of the approach. Simulation experiments using LTspice, a high-performance software developed by Linear Technology Corp, were conducted to gain a thorough understanding of the cell-attached system before performing real-world electrophysiological experiments.

During the simulations, pulses were delivered through the electrode, allowing us to measure the currents resulting from both the charge and discharge of a pulse. We also manipulated the charges between the capacitors to investigate their impact on the charge and discharge currents. The results demonstrated a strong correlation between the potential applied between the capacitors and the discharge current measurements. With sufficient data, machine learning algorithms can provide accurate predictions based on current measurements. This finding suggests great promise for accurately measuring cell membrane potential using the electrode-cell configuration without damaging the cells.

### Model Description

The bilayer structure of the cell membrane (11-14) can be likened to a capacitor in several ways. Like a capacitor, the cell membrane consists of two layers of lipid molecules, with hydrophilic heads facing outward and hydrophobic tails facing inward. This arrangement creates a separation of charge, similar to how a capacitor stores electrical charge. The lipid bilayer’s hydrophobic interior acts as an insulating barrier, preventing the free movement of ions and molecules across the membrane, much like the dielectric material in a capacitor. Additionally, the cell membrane exhibits capacitance-like behavior by allowing the selective movement of charged particles (such as ions) across the membrane, which is essential for electrical signaling and transport processes in cells.

Most cells carry a negative electric charge (Figure 1A). When an electrode is attached to the cell membrane in a bath solution, it can be viewed as two series-connected capacitors with a negative charge present inside (Figure 1B). In this arrangement, C1 represents the portion of the membrane where electrode is attached, while C2 corresponds to the remaining membrane. This analogy to capacitors in an electrical circuit allows for an understanding of the electrical properties exhibited by the cell membrane (Figure 1C).

**Figure 1.**
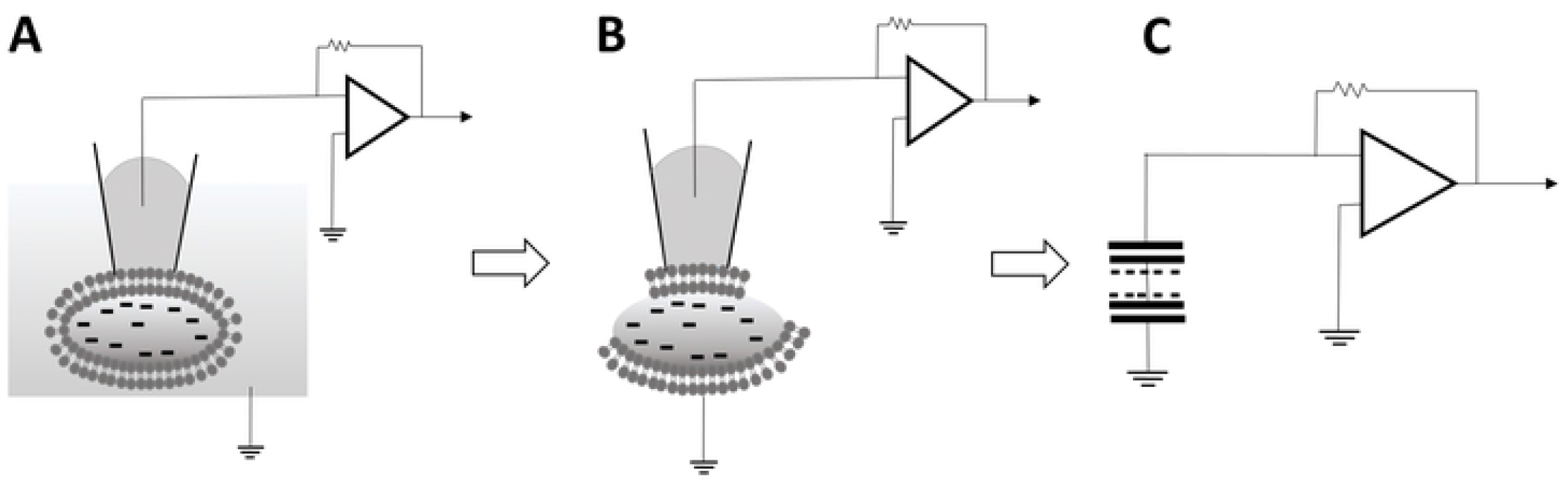
Model of Cell as Series-Connected Capacitors. (A) A cartoon depiction of cell-attached configuration is shown. (B) a visually representation of the two parts of the cell membrane, with one part held by the electrode and the other part representing the remaining membrane. (C) An illustrates the resemblance of this model to two capacitors with a negative charge situated in the middle.

### Figure 1: Model of Cell as Series-Connected Capacitors

For a circuit with two capacitors in series, the charge on each capacitor is the same, but the voltages will differ based on their capacitances. If V is the total applied voltage, the voltages across each capacitor, V1 and V2, are given by:

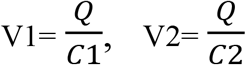

where V=V1+V2 and Q is the same for both capacitors.

If a negative charge Qneg is applied between the two series-connected capacitors, this charge will affect the voltage across each capacitor. Specifically:

When C1 is charged from the rising edge of a positive voltage pulse, the addition of a negative charge Qneg between the capacitors causes V1 to increase. The new voltage across C1 is given by:

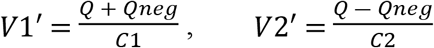

Conversely, when C1 is charged from the falling edge of a negative voltage pulse, the voltage V1 decreases due to the negative charge Qneg. The new voltage across C1 is given by:

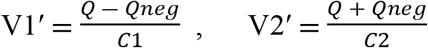

The changing voltage across the capacitors will, in turn, affect the charge and discharge current. The charge and discharge currents are given by the formula:

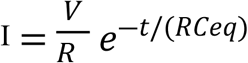

where V represents the voltage across the capacitors, R is the resistance in the circuit, *t* is time, and *RCeq* is the time constant determined by the resistance and series capacitance values. The charge and discharge current distribution is influenced by the voltage difference across each capacitor, regardless of the polarity of the applied pulse.

When a negative charge is introduced between the capacitors, it changes the voltage distribution across each capacitor (VC1, VC2). This change in voltage not only impacts the overall distribution of charge or discharge current but also influences the current distribution differently based on the polarity of the applied pulse (positive or negative). By carefully measuring and analyzing the resulting current distributions from both negative and positive pulses, valuable insights can be gained regarding the estimation of the magnitude of the negative charge present between the capacitors.

## Results

To test the theory, we simulated the model using LTspice (Version (x64): 24.0.9). The circuit diagram and component values are shown in Figures 2. 100 mV, 0.1 s test pulse (V1) was applied, and the potential (V2) between the two capacitors was varied from -100 mV to -10 mV in 10 mV increments to mimic the physiological charge range inside the cell. The current was measured from R1.

**Figure 2.**
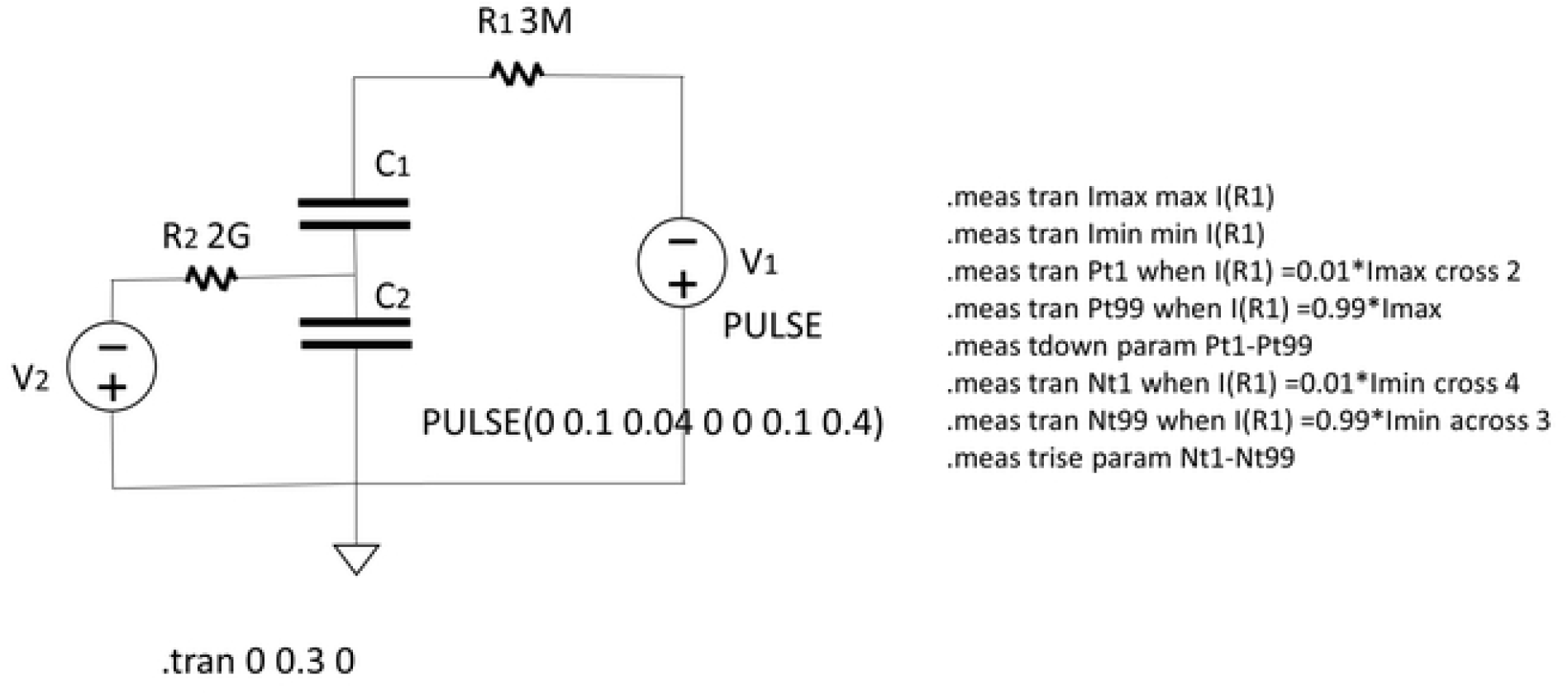
TLspice Circuit and Component Values. (A) the diagram of the simulation circuit in TLspice. (B) the values of the components used in the circuit. V1 represents the applied pulses to the circuit, a positive 100 mV 0.1 s pulse. V2 ranges from -10 mV to -110 mV, representing the internal potential of the cell.

### Figure 2: TLspice Circuit and Component Values

Since the charge and discharge currents follow the same underlying principles, the discharge current measurements obtained from a positive electric pulse are equivalent to the charge current measurements obtained from a negative pulse, assuming all other conditions remain constant. To compare positive and negative charges, we measured the charge and discharge currents at the rising and falling edges of a pulse.

We examined a total of 202 groups of capacitor combinations (S1). Typically, the electrode tip area (C1) was smaller compared to the remaining part of the cell membrane (C2). For C1, capacitor values ranged from 1 pF to 5.6 pF, and for C_2_, values ranged from 2.8 pF to 29.1 pF, with C2 always greater than C1. Electric pulses with an amplitude of 100 mV and a duration of 100 ms were used to ensure proper charging and discharging of the capacitors.

Figure 3 illustrates two distinct examples of capacitor combinations. In Figure 3A, the charge and discharge currents (represented by red curves) were generated by a 100 mV pulse (in blue, measured from the negative side of power source V1). The top panel shows C1 at 3.4 pF and C2 at 8.8 pF with V2 at -10 mV, while the bottom panel shows V2 at -60 mV. We measured the maximum positive current (Max, indicated by a green arrow) and minimum negative current (Min, indicated by a red arrow), along with the total charge (T1) and discharge time (T2). T1 and T2 refer to the duration between 99% and 1% of the maximum or minimum peak current, as indicated by the white and yellow dashed lines on the graph.

**Figure 3.**
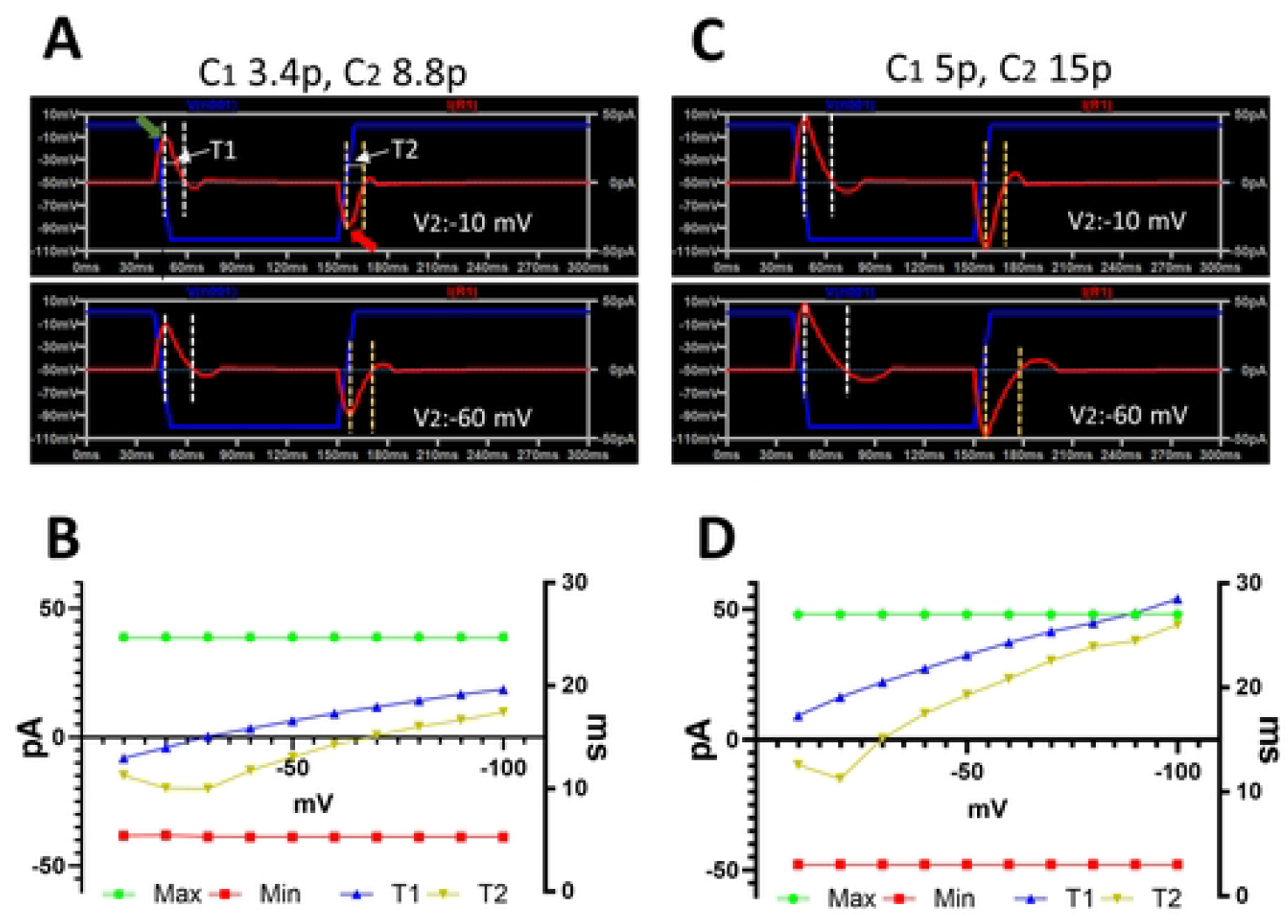
Charge and discharge Currents at Different Internal Potentials. (A) the charge and discharge currents (shown in red) are evoked by a 100 mV pulse (in green). The upper panel represents the currents when V2 is -10 mV, while the bottom panel represents the currents when V2 is -60 mV. The maximum and minimum currents (Max and Min) are indicated by green and red arrows respectively. The charge and discharge times are measured between 99% and 1% of the maximum (white dashed lines) or minimum (yellow dashed lines) currents. (B) Current measurements at varies V2 levels. C and D are another example for a different capacitor combination.

### Figure 3: Charge and discharge Currents at Different Internal Potentials

Figure 3B illustrates measurements taken at voltages (V2) ranging from -100 to -10 mV in 10 mV increments. The maximum (Max) and minimum (Min) currents, depicted by the green and red lines (left y-axis), show no significant changes with varying V2. However, the charge (T1) and discharge (T2) times, represented by the blue and yellow lines (right y-axis), increase as V2 becomes more negative.

Figures 3C and 3D provide another example with C1 at 5 pF and C_2_ at 15 pF. Similar to previous cases, as V2 becomes more negative, charge and discharge times increase, though the maximum and minimum currents do not change significantly.

Linear regression analysis revealed a strong correlation between V2 and the four measurements across all 202 groups of capacitor combinations, with R^2^ values ranging from 0.94 to 0.99. These consistent findings indicate that while the charge and discharge times are influenced by both the capacitors and the applied potential between them, the maximum and minimum currents are significantly affected only by different capacitor combinations. Additionally, the charge and discharge current distributions were affected differently by the varying conditions. These differences make it possible to predict the potential between the capacitors by measuring the currents.

To evaluate the prediction of V2 using charge and discharge current measurements, we employed the XGBRegressor machine learning model. The target variable, V2, ranged from -100 to -10 mV in 10 mV increments. We used four current measurement values (Max, Min, T1, and T2) as features and introduced two additional features: the ratio of maximum current to discharge time (Max/T2) and the difference between charge and discharge times (T1-T2) to enhance prediction accuracy. This resulted in a total sample size of 2020 with 6 features. We used 80% of the data for training and 20% for testing. The model achieved a mean squared error (MSE) of 80.28, an R^2^ score of 0.90, and a 5-fold cross-validated root mean squared error (RMSE) of 13.79 ± 3.30.

The MSE of 80.28 indicates a relatively low average difference between predicted and actual values. The R^2^ score of 0.90 suggests that our model explains 90% of the variance in the target variable, demonstrating a good fit to the data. The RMSE of 13.79 ± 3.30 reflects consistent model performance across different data folds, with a modest average error and low variability.

### Figure 4. Actual vs. Predicted Values and feature importance plot

Figure 4A shows the Actual vs. Predicted Values Plot, where data points align well along the diagonal line, particularly between 30 to 80 mV, indicating good model performance with some observed variation. Figure 4B highlights the feature importance plot, showing that the most significant feature is the time difference between total charge and discharge times (T1-T2). This suggests that the dynamics of how quickly the system charges and discharges play a crucial role in predicting the potential between the capacitors (V2).

**Figure 4.**
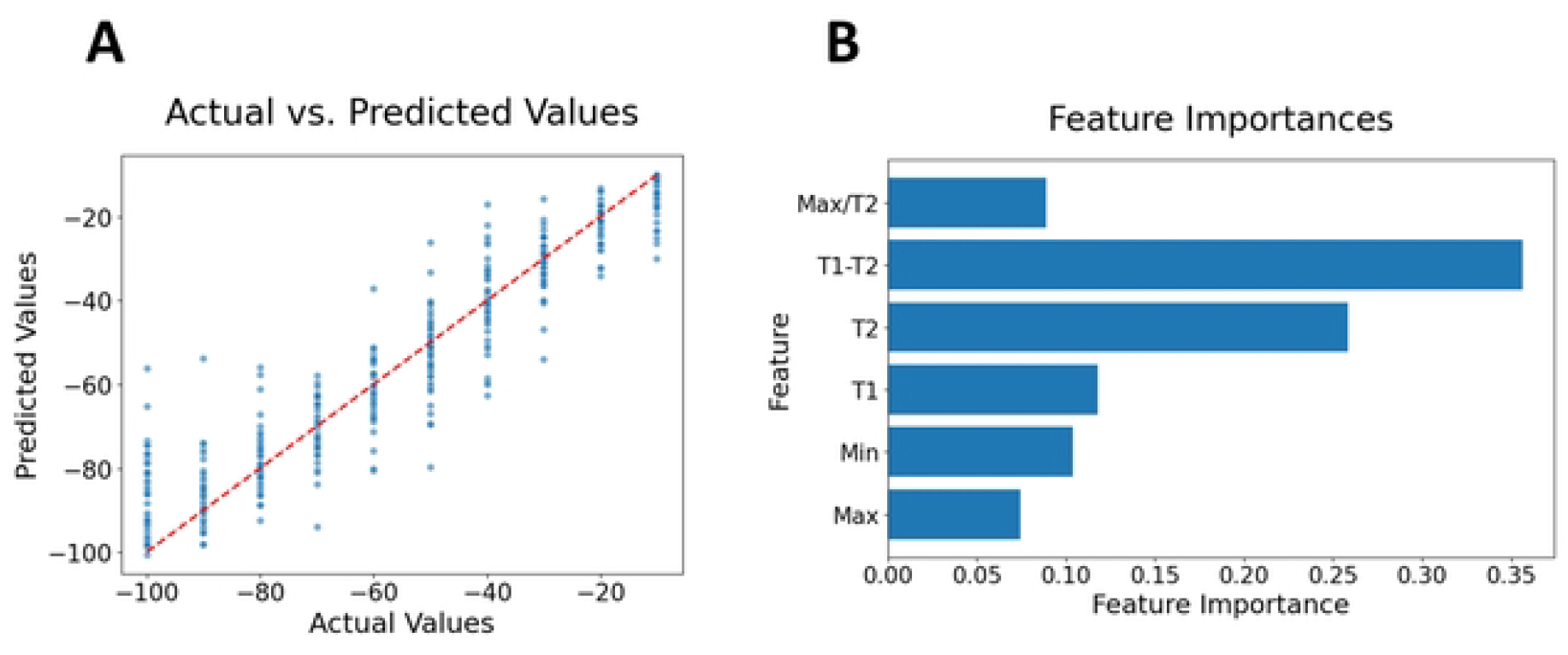
Actual vs. Predicted Values and feature importance plot. (A) Actual vs. Predicted Values Plot visually representing the alignment between the model’s predictions and the actual values. The data points align along the diagonal line, particularly between -30 to -80 mV. (B) The feature importance plot indicates that the most important feature is the time difference between total charge and discharge time.

For future experiments, refining the measurement of these key features could enhance prediction accuracy. This might involve improving the precision of time measurements and exploring additional parameters that could influence charge and discharge dynamics. The learning curve plot (not shown) suggests that increasing the sample size could further improve model accuracy, highlighting the importance of collecting more extensive data to capture system variability comprehensively.

## Discussion

This study, conducted through LTspice simulations, demonstrated that the negative charge between two capacitors can be accurately predicted by measuring the charge and discharge currents, even with a relatively small dataset. However, future experiments could enhance these predictions by increasing the data size and considering additional parameters such as the time constant, area under the curve, and charging slopes. Testing with different pulse characteristics could also improve accuracy and robustness.

Incorporating a DC filter into the experimental setup may provide benefits, such as reducing noise and mitigating artifacts from imperfect electrode seals, which could help in obtaining more reliable measurements.

Measuring action potentials from neurons with this method may be challenging due to the longer charge times used in this experiment compared to the brief duration of an action spike. To address this, combining extracellular recordings with electric pulse applications could provide the necessary temporal resolution to capture rapid changes in membrane potential.

Validating these results with real cells is essential. Biological experiments are inherently complex due to the intricate internal structure of cells and the dynamic nature of biological systems. The whole-cell patch clamp technique, which allows precise control of internal potential and accurate measurement of discharge currents, is crucial for this validation. Additionally, voltage-sensitive dyes can be valuable in validating the predictions. By employing these techniques, we can thoroughly investigate the relationship between membrane potential and current characteristics.

The whole-cell patch clamp and voltage-sensitive dye techniques enable detailed investigations into how membrane potential affects charge and discharge currents. By collecting comprehensive experimental data and applying machine learning algorithms, we can establish a strong correlation between these currents and membrane potential, offering a reliable, non-invasive method for assessing membrane potential.

This non-invasive approach has significant potential for future research into cell membrane dynamics and drug discovery. Integrating insights from simulations and experimental validations will deepen our understanding of membrane potential effects, confirming the capability to measure membrane potential accurately without damaging cells.

## Reference

1. Hille B, 2001. Ion Channels of Excitable Membranes (3rd ed.). Sinauer Associates.

2. Khadria A. 2022. Tools to measure membrane potential of neurons. Biomed J., 45(5):749–762.

3. H Antonická, H. D Floryk, P Klement, L Stratilová, J Hermanská, H Houstková, M Kalous, Z Drahota, J Zeman and J Houstek. 1999, Defective kinetics of cytochrome c oxidase and alteration of mitochondrial membrane potential in fibroblasts and cytoplasmic hybrid cells with the mutation for myoclonus epilepsy with ragged-red fibres (‘MERRF’) at position 8344 nt. Biochem J. 15;342 Pt 3(Pt 3):537–44.

4. M Hiraoka, S Kawano, Y Hirano and T Furukawa. 1998. Role of cardiac chloride currents in changes in action potential characteristics and arrhythmias. Cardiovasc Res. 40(1):23–33.

5. Y Jammes, N Adjriou, N Kipson, C Criado, C Charpin, S Rebaudet, C Stavris, R Guieu, E Fenouillet and F Retornaz. 2020. Altered muscle membrane potential and redox status differentiates two subgroups of patients with chronic fatigue syndrome. J Transl Med. 19;18(1):173–183.

6. GJ. Kaczorowski, OB. McManus, BT. Priest and ML. Garcia. 2008. Ion Channels as Drug Targets: The Next GPCRs. J Gen Physiol. 131(5): 399–405.

7. Fluhler E, Burnham VG and Loew LM. Spectra membrane binding, and potentiometric responses of new charge shift probes. 1985. Biochemistry. 24: 5749–5755.

8. HK Zhang, P Yan, J Kang, DS Abou, HND Le, AK Jha, DLJ Thorek, JU Kang, A Rahmim, DF Wong, et al. 2017. Listening to membrane potential: photoacoustic voltage-sensitive dye recording. Journal of Biomedical Optics. 22: 45006.

9. Peterka DS, Takahashi H and Yuste R. 2011. Imaging voltage in neurons. Neuron, 69(1), 9–21.

10. JQ Xu, YL Liu, Q Wang, HH Duo, XW Zhang, YT Li and WH Huang. 2015. Photocatalytically Renewable Micro-electrochemical Sensor for Real-Time Monitoring of Cells. Angewandte Chemie International Edition. 54: 14402–14406.

11. Singer SJ, and Nicolson GL. 1972. The Fluid Mosaic Model of the Structure of Cell Membranes. Science. 175: 720–731.

12. Ingólfsson HI, Melo MN and Roux B. 2014. Lipid Organization of the Plasma Membrane. Journal of the American Chemical Society. 136(41), 14554–14559.

13. Tanaka M, Sackmann E. 2005. Polymer-supported membranes as models of the cell surface. Nature 437:656–663.

14. T Róg, M Pasenkiewicz-Gierula, I Vattulainen and M Karttunen. 2009. Ordering effects of cholesterol and its analogues. Biochimica et Biophysica Acta (BBA) - Biomembranes, 1788(1), 97–121.

15. Ueda Y, Sato M. 2018. Cell membrane dynamics induction using optogenetic tools Biochem Biophys Res Commun. 25;506(2):387–393.

16. Bezanilla F. 2018. Gating currents J Gen Physiol. 2;150(7):911–932.

